# Suppressor mutations in *Mecp2*-null mice reveal that the DNA damage response is key to Rett syndrome pathology

**DOI:** 10.1101/810929

**Authors:** Adebola Enikanolaiye, Julie Ruston, Rong Zeng, Christine Taylor, Marijke Shrock, Christie M. Buchovecky, Jay Shendure, Elif Acar, Monica J. Justice

## Abstract

Mutations in X-linked methyl-CpG-binding protein 2 (*MECP2)* cause Rett syndrome (RTT). We carried out a genetic screen for secondary mutations that improved phenotypes in *Mecp2*/Y mice after mutagenesis with *N*-ethyl-*N*-nitrosourea (ENU), aiming to identify potential therapeutic entry points. Here we report the isolation of 106 founder animals that show suppression of *Mecp2*-null traits from screening 3,177 *Mecp2*/Y genomes. Using exome sequencing, genetic crosses and association analysis, we identify 33 candidate genes in 30 of the suppressor lines. A network analysis shows that 61% of the candidate genes cluster into the functional categories of transcriptional repression, chromatin modification or DNA repair, delineating a pathway relationship with MECP2. Many mutations lie in genes that are predicted to modulate synaptic signaling or lipid homeostasis. Surprisingly, mutations in genes that function in the DNA damage response (DDR) also improve symptoms in *Mecp2/Y* mice. The combinatorial effects of multiple loci can be resolved by employing association analysis. One line, which was previously reported to carry a suppressor mutation in a gene required for cholesterol synthesis, *Sqle*, carries a second mutation in retinoblastoma binding protein 8 (*Rbbp8* or *CtIP*), which regulates a DDR choice in double stranded break (DSB) repair. Cells from *Mecp2*/Y mice have increased DSBs, so this finding suggests that the balance between homology directed repair and non-homologous end joining is important for neuronal cells. In this and other lines, the presence of two suppressor mutations confers better symptom improvement than one locus alone, suggesting that combination therapies could be effective in RTT.

## Introduction

Mouse genetics is a powerful tool to identify molecular mechanisms that are important for disease suppression. Employing a modifier screen, a dominant mutation can be isolated by its ability to alter the presentation of a recessive or dominant trait to discover genes that act in a given developmental or biochemical pathway. Modifier screens have been rare in the mouse; however, using NextGen sequencing technologies, candidate mutations can now be efficiently identified without extensive breeding and mapping. Mutagenesis screens that focus on disease suppression may identify unrecognized pathways of pathogenesis as alternative therapeutic entry points.

Rett syndrome (RTT) is a prototype disease for which a modifier screen would be beneficial and representative for other diseases. RTT is a nearly monogenic disorder with over 95% of patients carrying mutations in methyl-CpG-binding protein 2 (*MECP2*), a gene not present in invertebrates. MECP2 is central to neurological function and is associated with other diseases, including intellectual disabilities, autism, neuropsychiatric disorders and lupus erythematosis (Neul and Zoghbi 2004; Bienvenu and Chelly 2006). The type of *MECP2* mutation does not always correlate with disease severity, in part, due to skewing of X-chromosome inactivation in heterozygous females (Shahbazian et al. 2002). Hemizygous males with truncating or loss of function mutations in *MECP2* have a more severe phenotype than females with RTT and usually die by two years of age (Bienvenu and Chelly 2006).

Mouse models recapitulate most of the symptoms of RTT and are crucial for understanding why symptoms arise (Chen et al. 2001; Guy et al. 2001). Male mice that lack *Mecp2* are normal through three weeks of age, but they develop hypoactivity, limb clasping, tremors, and abnormal breathing as early as four weeks, depending upon the allele and the genetic background (Katz et al. 2012). The symptoms become progressively worse, leading to their death at 6 – 12 weeks. RTT has historically been considered a neurological disease (Jellinger and Seitelberger 1986). MECP2 is expressed at near histone levels in neurons, and neurons of *Mecp2*/Y mice exhibit a number of deficits, including delayed transition into mature stages, altered expression of presynaptic proteins and reduced dendritic spine density (Smrt et al. 2007; Belichenko et al. 2009). Since dendritic spines are the locations of excitatory synapses, a consequence of *Mecp2* deficiency is reduced synaptic number and functional connectivity (Chao et al. 2007; Chapleau et al. 2009). However, MECP2’s expression outside the nervous system leads to a number of systemic problems, including metabolic syndrome (Kyle et al. 2016; Ross et al. 2016; Kyle et al. 2018).

Multi-allelic contributions or environmental effects may cause the wide degree of phenotypic variation that is common to many human diseases (Enikanolaiye and Justice 2019). Rare familial cases of RTT suggest that genetic modification can alter the severity of symptoms (Ravn et al. 2011; Grillo et al. 2012), but as RTT is a rare disorder, finding the molecular basis for phenotypic modification in humans is daunting. Identifying genes that when mutated ameliorate or prevent progression of symptoms in a mouse model would help to focus efforts towards understanding the functions of MECP2 and how the diseases ensues upon its loss. An unbiased modifier screen, which is intended to alter the phenotypic outcome, may reveal genes that act in common developmental or biochemical pathways with MECP2.

Further, identifying the genes that improve the RTT-like phenotype in mice may allow for the discovery of pathways that inform therapeutic strategies in humans. In a preliminary dominant suppressor mutagenesis screen using the supermutagen N-ethyl-N-nitrosourea (ENU), we isolated five modifier loci, named Suppressor of Mecp2 (*Sum) 1 – 5*, which suppress or ameliorate the symptoms of *Mecp2* mutation in male mice (Buchovecky et al. 2013). We previously reported that the lesion responsible for disease amelioration at the *Sum3^m1Jus^* locus is a stop codon mutation in a rate-limiting enzyme in cholesterol biosynthesis, squalene epoxidase (*Sqle*). This finding led to the discovery that cholesterol and lipid homeostasis is perturbed in *Mecp2*-null mice, and that statin drugs ameliorate symptoms as well (Buchovecky et al. 2013). This discovery suggested that finding additional modifiers of *Mecp2-*null phenotypes would lead to a better understanding of *Mecp2’*s functions and how its mutation could lead to pathological changes, while unmasking new therapeutic options for RTT.

Our overarching goal was to identify a large number of suppressors such that a list of potential therapeutic targets could be developed for RTT. Here we report the screening of an additional 2,498 gametes to total 3,177 gametes screened for suppressor mutations. In this second screen, 96 additional lines that show evidence of improving a variety of subjective health traits were isolated. Next generation whole exome sequencing has identified over 3,600 potentially causative lesions in 72 of these lines. We have solved the genetic cause in 13 lines to identify lesions in 23 genes that primarily fall into three pathways that contribute to RTT-like symptoms in mice. Altogether, lesions in alleles of solved genes or pathway components identify a possible cause of symptom improvement in 30 of 106 lines that had improved traits.

## Results

### Two suppressor screens produce 106 founders with trait improvement

ENU-treated C57BL/6J males were bred to female 129.*Mecp2*^tm1.1Bird^/+ mice (Figure 1A). Male offspring in the first generation (N1), asymptomatic at weaning, were genotyped for presence of mutant *Mecp2*, and examined for amelioration of disease phenotypes by a dominant mutation. The preliminary screen 1, which was carried out in 2007 – 2009, totaled 2,963 animals of which 679 were *Mecp2*/Y (Figure 1A, gray) (Buchovecky et al. 2013). This number is close to one genome’s equivalent, since the mutation rate for the 3 × 100 mg/kg dose of ENU is about 1 new mutation per gene in every 655 gametes screened (Hitotsumachi et al. 1985). In screen 1, 10 lines showing suppression of traits were identified, but only six of these bred naturally and one (1527) was reconstituted by IVF of fresh sperm, because he did not show interest in mating. A non-standard five point scoring system was used to individually assess limb clasping, tremors, body weight and activity. In general, improvement of these scores indicated improved longevity as evidenced by the long lifetime of the founders and their offspring (Buchovecky et al. 2013).

**Figure 1:**
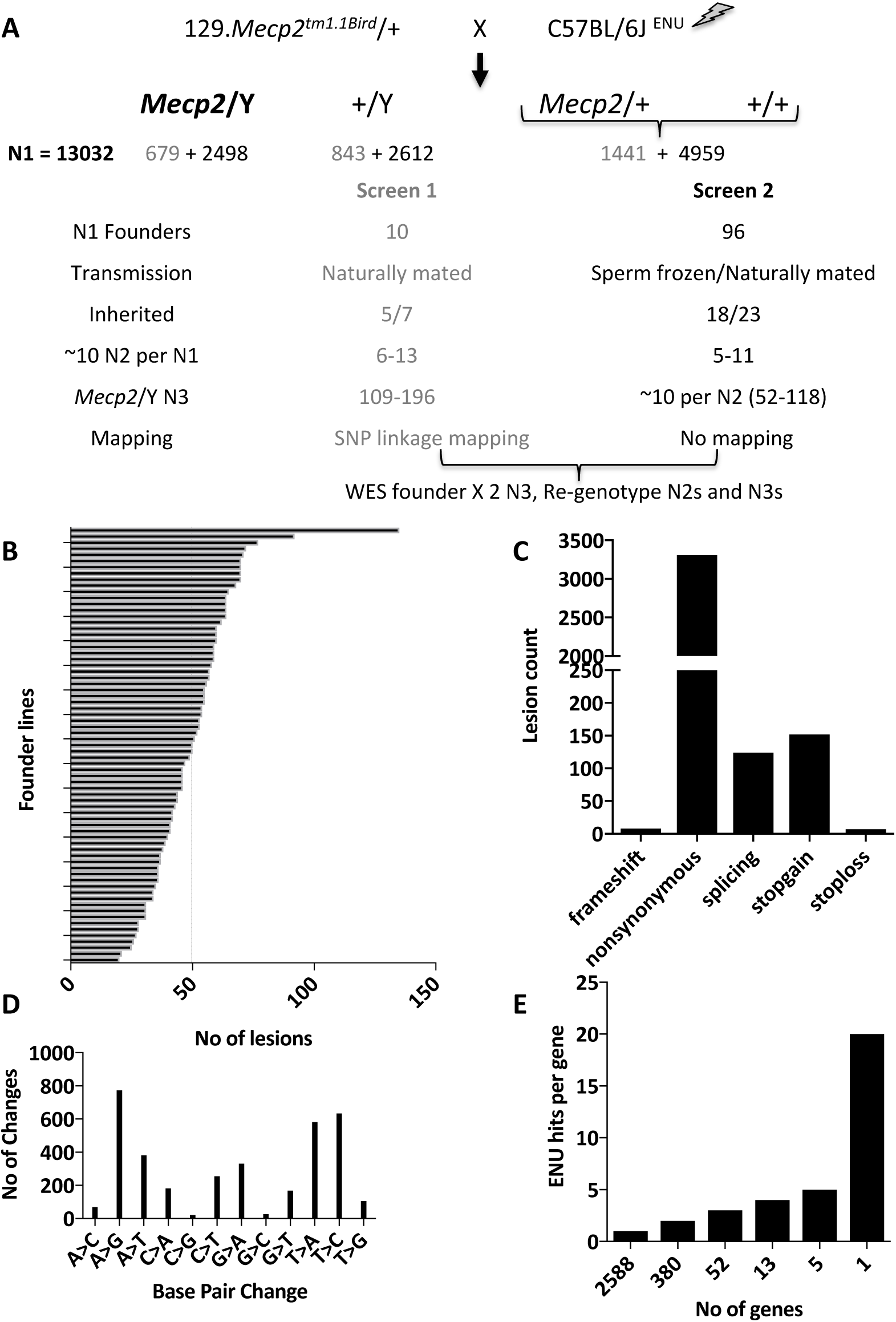
Overview of two dominant suppressor ENU mutagenesis screens in the *Mecp2^tm1.1Bird^* mouse model of RTT. A. Male C57BL/6J male mice were treated with ENU and mated to 129.*Mecp2^tm1.1Bird^*/+ females. Numbers obtained from Screen 1, already published, are in gray, and Screen 2 in black font. A subset of N1 male founder mice carrying the mutant *Mecp2* allele and showing suppression of disease phenotypes were bred (naturally or by IVF) for inheritance. N2 offspring were mated to 129.*Mecp2^tm1.1Bird^*/+ females or 129S6/SvEvTac males to generate N3s, of which two animals from two different N2 families that showed trait improvement were then whole exome sequenced (WES), and compared to the founder sequence. N3 offspring were genotyped to analyze for linkage or association. B. The WES was analyzed for 72 founder lines. The number of potentially causative lesions in each founder ranged from 20 to 135, with a mean of 50. C. The types of ENU-induced lesions generated were mostly nonsynonymous missense mutations (92%), followed by nonsense mutations (4%) and splicing mutations (3%). Frameshift mutations or the loss of a STOP codon accounted for the remainder (1%). D. The most common types of mutations induced by ENU were A to G and T to C transition mutations (22% and 18% respectively) while the fewest were C to G and G to C transversions (both less than 1%). E. The majority of genes with new ENU-induced lesions were unique, with 2588 genes carrying only one mutation.

In screen 2, carried out in 2013-2017, an additional 10,069 animals of which 2,498 were *Mecp2*/Y produced 96 additional N1 founder males that showed suppression of one or more traits. In screen 2, sperm from each founder was frozen, and of 23 lines mated, 9 bred naturally before they were sacrificed whereas 14 were reconstituted using IVF of frozen sperm. In screen 2, all N1 mice were evaluated weekly starting at P35 for four subjective health parameters that have previously been noted in *Mecp2/Y* mice (Guy et al. 2001): hindlimb clasping, tremors, activity and general body condition. Muscle tone was identified as an important indicator of health and was also assessed. Each subjective parameter was scored out of 2, whereby 0 indicated the health trait was equivalent to wild type, 1 if intermediate, or 2 if severe. Body weights were also obtained weekly and were scored 0 if less than (<) 29 grams (g), 1 if ≥ 30-34g, and 2 if over (>) 35g. These five subjective health parameters and body weight were totaled to obtain a score out of 12. In screen 2, animals were considered to be improved if their summed health score totaled < 8 at eight weeks of age, and they were maintained only until 12 - 14 weeks, when they were sacrificed for sperm cryopreservation. N3 animals were evaluated using the same criteria. Of 23 families mated in screen 2, 18 showed evidence of inherited suppression of one or more traits, but five did not show evidence of inheritance. Notably, the *Mecp2*/Y animals from the strains maintained at the two institutions had different life spans. Therefore, *Mecp2*/Y animals from screen 1 were considered improved if they lived ≥ 14 weeks, while those from screen 2 were considered improved if they lived ≥ 10 weeks and had improved scores on assessed health traits.

### Whole exome sequencing identifies candidate genes in founder males

When the first modifiers were isolated in screen 1, whole exome sequencing (WES) was not yet available, so the first line was sequenced by custom capture re-sequencing of a mapped region (line 352). WES was carried out on lines 856, 895, 1395 and 1527, but the sequence was analyzed only in the regions with positive LOD scores obtained by mapping, and only one candidate gene, *Sqle*, from line 895 was reported (Buchovecky et al. 2013). The WES of 69 of the founder N1 males from the second screen was analyzed for all potential contributing mutations throughout the genome (Supplemental Table S1). Naturally-occurring single nucleotide polymorphisms (SNPs) that were specific to the 129S6/SvEv and C57BL/6J strains were filtered by comparing with parental and all sequenced founder male genomes. New ENU-induced mutations would represent rare variants, and would most frequently occur as single base pair changes and rarely, small frameshifts or deletions. Many ENU-induced variants are not detrimental, so variants that are predicted to be tolerated were excluded (Miosge et al. 2015). Potentially causative variants found in each N1 founder male ranged in number from 20 to 135, with a mean of 50 (Figure 1B), which is consistent with numbers found in other ENU screens (Wang et al. 2015). The 3,601 potentially causative variants were largely nonsynonymous missense mutations (3310, 92%), 124 (3%) were splicing mutations, 152 (4%) were nonsense mutations, and others were either frameshift or loss of a STOP codon (1%) (Figure 1C). Most of the lesions were transition mutations (A to G (22%) and T to C (18%)), closely followed by transversions (A to T (11%) and T to A (16%)) (Figure 1D), consistent with the most common types of base changes found after germline mutagenesis with ENU (Justice et al. 1999). Each of 2588 genes had only one potentially causative lesion found in the N1 males, each of 380 genes had two different lesions, 52 genes had three alleles, 13 genes had four, five genes had five, and a single large gene, Titan (*Ttn*), had 20 alleles (Figure 1E).

### Confirming the candidate lesions shows complex inheritance of traits

The inheritance of traits was determined from the results of mating 30 N1 founders from both screens (7 from screen 1 and 23 from screen 2) to 129.*Mecp2/+* females. Their N2 female offspring, carrying the *Mecp2* mutation inherited from their father, were mated again to 129S6/SvEvTac males, whereas N2 male offspring were mated to 129.*Mecp2/+* females. In screen 1, each line was mapped to narrow the suppressor locus to a chromosomal location using SNP panels prior to sequencing (Neuhaus and Beier 1998; Moran et al. 2006). In screen 2, SNPs were not employed for mapping. When possible, up to 10 N2 offspring from each family were mated to produce approximately 10 *Mecp2*/Y N3 offspring, which were assessed for phenotype and aged to at least 12 weeks. If the modifier was inherited in a Mendelian fashion, 50% of the N2 animals should have inherited the trait and 50% then should pass it on to their N3 offspring; thus, only 25% of the total N3 offspring are expected to inherit the suppressor trait (Table 1). Of the lines in which the offspring were analyzed for suppression of traits, 23 showed evidence for inheritance of disease improvement (Supplemental Table S2). Moreover, all but four of the 23 lines segregated modifier loci in a non-Mendelian fashion, suggesting involvement of more than one locus (Table 1).

**Table 1:** Assessment of Mendelian inheritance of suppressor loci

| Line | Families with improvement |  | # N3 Mecp/Y Improved/ Total |  | Inheritance (p-val) | Solved |
| --- | --- | --- | --- | --- | --- | --- |
| 352* | 6/6 | 1.00 | 36/129 | 0.28 | <0.001 | *Prdm15*<sup>Sum1 m1Jus</sup> |
| 856* | 10/10 | 1.00 | 76/165 | 0.46 | 0.003 | *Zzz3*<sup>Sum2 m1Jus</sup><br>Chr 6: LOD 3.9 |
| 895* | 8/9 | 0.89 | 32/109 | 0.29 | <0.001 | *Sqle*<sup>Sum3 m1Jus</sup><br>*Rbbp8*<sup>Sum6 m1Jus</sup> |
| 520 | 8/8 | 1.00 | 19/116 | 0.16 | <0.001 | *CD22*<sup>Sum7 m1Jus</sup><br>*Clint1*<sup>Sum7i</sup><br>*Kdm4a*<sup>Sum8 m1Jus</sup> |
| 4654* | 5/9 | 0.56 | 8/81 | 0.10 | <0.001 | *Tm7sf2*<sup>Sum9 m1Jus</sup><br>*Dennd4a*<sup>Sum9i</sup><br>*Gtf3c5*<sup>Sum10 m1Jus</sup> |
| 4751* | 5/6 | 0.83 | 13/81 | 0.16 | <0.001 | *Arhgef15*<sup>Sum11 m1Jus</sup><br>*Spin1*<sup>Sum12 m1Jus</sup> |
| 4799* | 4/7 | 0.57 | 8/70 | 0.11 | <0.001 | *Aacs*<sup>Sum13 m1Jus</sup><br>*Atp8a1*<sup>Sum14 m1Jus</sup> |
| 918-15L* | 3/6 | 0.50 | 7/60 | 0.12 | <0.001 | *Celsr3*<sup>Sum15 m1Jus</sup> |
| J_57L | 7/9 | 0.78 | 18/108 | 0.17 | <0.001 | *Hcn2*<sup>Sum16 m1Jus</sup><br>*Apoa5*<sup>Sum17 m1Jus</sup><br>*Col24a1*<sup>Sum17i</sup> |
| A_27R | 3/10 | 0.30 | 8/103 | 0.08 | <0.001 | *Tet1*<sup>Sum18 m1Jus</sup> |
| A_333N | 3/8 | 0.38 | 9/62 | 0.15 | 0.009 | *Fan1*<sup>Sum19 m1Jus</sup> |
| M_199LL* | 4/5 | 0.80 | 9/52 | 0.17 | 0.002 | *Birc6*<sup>Sum20 m1Jus</sup><br>*Dbnl*<sup>Sum20i</sup> |
| M_333R* | 5/6 | 0.83 | 10/88 | 0.11 | <0.001 | *Zbtb41*<sup>Sum21m1Jus</sup> |
| 1395* | 6/6 | 1.00 | 70/196 | 0.36 | <0.001 | Chr 7: *Sum4*<sup>m1Jus</sup> |
| 1527 | 13/14 | 0.93 | 61/190 | 0.32 | <0.001 | *Sum5*<sup>m1Jus</sup> |
| 136* | 7/11 | 0.64 | 12/104 | 0.12 | <0.001 | *Tet1*<sup>Sum18 m2Jus</sup> |
| 137* | 7/10 | 0.70 | 7/101 | 0.07 | <0.001 |  |
| 591 | 7/9 | 0.78 | 7/114 | 0.06 | <0.001 | *Tet1*<sup>Sum18 m3Jus</sup> |
| 722 | 4/10 | 0.40 | 8/102 | 0.08 | <0.001 |  |
| 933-15N | 4/10 | 0.40 | 5/118 | 0.04 | <0.001 |  |
| A_87N | 4/9 | 0.44 | 8/95 | 0.08 | <0.001 |  |
| A_120N | 3/7 | 0.43 | 8/70 | 0.11 | 0.002 |  |
| A_134L* | 3/5 | 0.60 | 8/84 | 0.10 | <0.001 |  |
| A_230RR | NM |  |  |  |  | *Arhgef15*<sup>Sum11 m2Jus</sup> |
| 604 | NM |  |  |  |  | *Celsr3*<sup>Sum15 m2Jus</sup> |
| A_347N | NM |  |  |  |  | *Celsr3*<sup>Sum15 m3Jus</sup> |
| A_83L | NM |  |  |  |  | *Birc6*<sup>Sum20 m2Jus</sup> |
| 620 | NM |  |  |  |  | *Birc6*<sup>Sum20 m3Jus</sup><br>*Ncor1*<br>*Brca1* |
| S_165N | NM |  |  |  |  | *Ncor1* |
| A_189N | NM |  |  |  |  | *Brca1* |
| J_157N | NM |  |  |  |  | *Tbl1xr1* |
| J_136N | NM |  |  |  |  | *Zbtb41*<sup>Sum21 m2Jus</sup> |
| M_420L | NM |  |  |  |  | *Mre11a* |
| M_79N | NM |  |  |  |  | *Rad50* |
| A_173N | NM |  |  |  |  | *Brca2* |
| 920_39R | NM |  |  |  |  | *Cyp46a1* |
| 4633 | NM |  |  |  |  | *Rcor1* |
| 272 | NM |  |  |  |  | *Sin3a* |

\* =Line mated naturally, NM= not mated

In four lines, 352, 856, 520 and 1395, all N2 families showed at least one animal with a degree of improvement. In line 856, the pattern of inheritance suggested that either locus conferred improvement, while the inheritance of both loci suggested additive health improvement; in contrast, in line 520 no animals showed greatly improved health scores, suggesting that the two loci may have influenced health traits independently. Seven lines, 137, 591, 722, 933-15N, A_87N, A_120N and A_134L, were not solved because so few *Mecp2*/Y N3 animals showed trait improvement (fewer than 8 total animals per line), possibly because two or more loci must be inherited together to see an effect. In all cases, because these are complex loci, it is also possible that a genetic combination resulted in decreased health or longevity, rather than improvement.

To identify candidate genes in screen 2, the exome from a minimum of two N3 offspring from different N2 parents was also sequenced. In each line, the N3 animals that exhibited the largest degree of improvement and belonged to families that had multiple animals with improvement were chosen for sequencing. Lesions that occurred in the N1 founder and these two offspring were considered as causative. Common SNPs identified in WES were used to genotype N3 animals to confirm the location after candidate loci were identified. Individual members of families that inherited trait improvement were re-genotyped for lesions that were potentially causative (Figure 1A; Supplemental Table S2).

Statistical analysis of candidate genes shows support for a solution in 13/23 lines (Table 1, 2; Supplemental Table S3). Within these 13 lines, 23 genes are supported by association analysis as candidates for suppression of a variety of traits. Interestingly, eight additional lines carry alleles of these 23 candidate genes, and 11 lines carry mutations in a related pathway member or *Mecp2* co-repressor complex member. Some lines carry multiple causative mutations that are supported, thus, in total candidate genes have been identified in 30 lines (Table 1, 2). The new loci that are supported by mating and association analysis are named *Sum 6 – 21*, since the first five loci were named in screen 1 (Buchovecky et al. 2013) (Table 1). Candidate genes are now identified for two of the lines from screen 1 (*Prdm15^Sum1m1Jus^* and *Zzz3^Sum2m1Jus^*). A second suppressor was found in line 895 from screen 1 (*Rbbp8^Sum6m1Jus^*). Loci that produced an association score only in the presence of another supported suppressor locus are named *Sum7i* (*Clint1*), *Sum9i* (*Dennd4a*), *Sum17i* (*Col24a1*), and *Sum20i* (*Dbnl*) for the interaction with that specific locus.

### Candidate genes fall into similar functional pathways

Genes with supported or pathway lesions were integrated into biological networks using the publicly available tool Cytoscape (v3.7) (Shannon et al. 2003) and its accompanying applications GeneMANIA (v3.5.1) (Warde-Farley et al. 2010) and ClusterMaker (1.3.1) (Morris et al. 2011). A functional enrichment map (network) determined by co-expression, shared protein domains, physical interactions or predictions revealed a total of 113 links (edges) amongst the proteins encoded by the candidate genes isolated in this screen (Figure 2). This network was further clustered into biological pathways confirming that most of the genes fall into three major pathways; lipid metabolism and homeostasis, synaptic function, and DNA damage response, with *Mecp2, Ncor1, Rcor1, Sin3a, Tbl1xr1* and *Rbbp8* overlapping with the transcriptional repression pathway. Gene ontology (GO) analysis revealed that the top 15 enriched terms in this network, as determined by their false discovery rate (FDR)-adjusted p-values, included previously documented MECP2-associated biological processes involved in transcriptional co-repressor activity and chromatin-associated functions. However, it also revealed novel GO terms such as DNA recombination, double-strand break repair and regulation of DNA metabolic processes. Thus 20/33 or 61% of the genes cluster into the general category of “Regulation of DNA Activity”, which might be expected for a functional relationship with MECP2. The genes with unknown function (*Clint1, Col24a1, Dennd4a*) fall outside the functional pathways.

**Figure 2:**
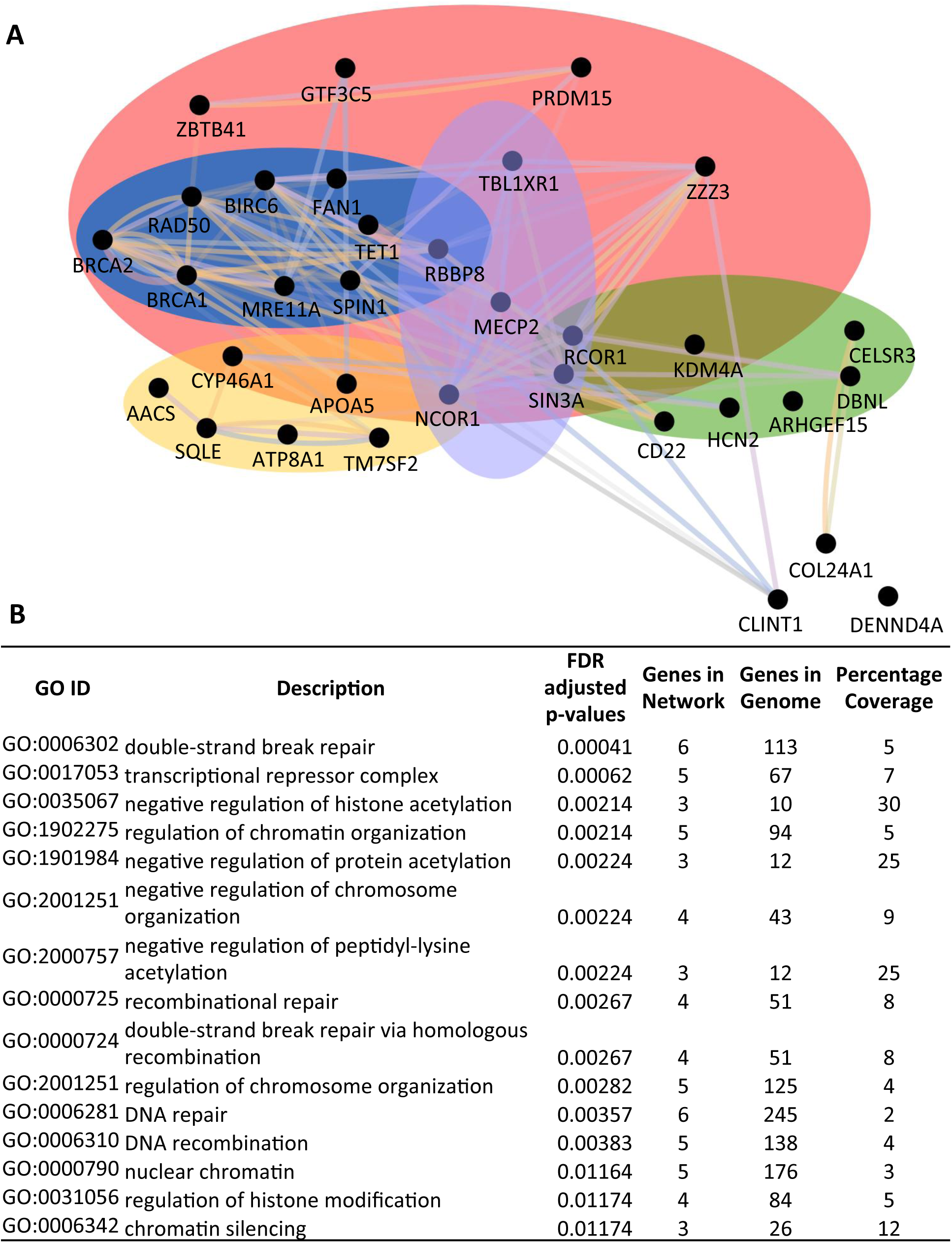
A functional network analysis of proteins encoded by candidate genes. A. A network enrichment map showing the relationships between proteins with supported or pathway lesions. Proteins are represented as black dots or nodes and the relationships between them are represented as colored lines or edges where purple lines denote co-expression, pink lines denote physical interactions, orange lines denote predicted interactions, blue lines denote co-localization and green lines denote shared protein domains. Further clustering (indicated by colored ovals) revealed that most of the proteins group into the broad categories of lipid metabolism and homeostasis (yellow), synaptic function (green), and DNA damage response (blue). Several proteins are also involved in transcriptional repression (purple). Sixty-one percent are included in the broad category regulation of DNA activity (pink). B. The most enriched gene ontology (GO) terms in this network, as determined by their false discovery rates (FDR)-adjusted P-value, are double-strand break repair, transcriptional repressor complex, negative regulation of histone acetylation and regulation of chromatin organization. The percentage coverage indicates the proportion of genes within a given GO ID that are also present in the network.

The co-repressors *Ncor1,* transducin (beta)-like 1X-linked receptor 1(*Tbl1xr1*), *Rcor1* and *Sin3a* are transcriptional repressors that act with MECP2, yet missense mutations in these genes occurred in lines that were not mated, so they are included in Table 2 but are not given *Sum* designations. Several additional supported genes are particularly intriguing because they are predicted to alter chromatin structure: these include PR domain containing 15 (*Prdm15*), zinc finger ZZ domain containing 3 (*Zzz3*), and lysine (K)-specific demethylase 4A (*Kdm4a*) (Shanle et al. 2017). The PR domain proteins, many of which have histone methyltransferase activity and others that recruit methyltransferases to DNA, play roles as molecular switches in many developmental processes (Fog et al. 2012; Hohenauer and Moore 2012). Little is known about PRDM15, but it functions in pluripotency (Mzoughi et al. 2017) and maps to the Down syndrome trisomy region of mouse Chromosome 16 and human Chromosome 21. ZZZ3 is a part of the Ada-Two-A-containing (ATAC) histone acetyltransferase complex, which widely regulates gene expression (Mi et al. 2018). KDM4A is important for the structure of heterochromatin during embryonic development, and also influences neuropathic pain through brain derived neurotrophic factor (BDNF) expression (Sankar et al. 2017; Zhou et al. 2017).

**Table 2:** Mutations in modifier lines

| Gene, Chromosome | Supported Lesion | Improvement p<0.5 | Allele or Pathway Lesion |
| --- | --- | --- | --- |
| **DNA damage response** |  |  |  |
| *Birc6*, Chr 17, NM_007566 | exon49:c.9611G>A:p.R3204Q (M_199LL) | health, weight<br>***Birc6*↔*Dbnl* Longevity** | exon23:c.G4784A:p.S1595N(A_83L)<br>exon66:c.G13516A:p.A4506T (620) |
| *Brca1*, Chr 11, NM_009764 |  |  | exon10:c.T679A:p.F227I (620)<br>exon4:c.A148G:p.K50E (A_189N) |
| *Brca2*, Chr 5, NM_001081001 |  |  | exon11:c.G2107T:p.D703Y (A_173N) |
| *Fan1*, Chr 7, NM_177893 | exon10:c.2392T>C:p.S798P (A_333) | Longevity, limb clasping, health |  |
| *Mre11a*, Chr 9, NM_018736 |  |  | exon17:c.T1915A:p.Y639N (M_420L) |
| *Rad50*, Chr 11, NM_009012 |  |  | exon16:c.A2717G:p.K906R (M_79N) |
| *Rbbp8*, Chr 18, NM_001081223 | exon15:c.2227C>T:p.Q743X (895) | Longevity<br>***Rbbp8*↔*Sqle* Longevity** |  |
| *Spin1*, Chr 13, NM_146043 | exon4:c.696C>A:p.Y232X (4751) | Longevity, limb clasping |  |
| *Tet1*, Chr 10, NM_027384 | exon1:c.241A>T:p.M81L (A_27R) | Longevity, muscle tone, activity | exon2:c.G1909T:p.V637F (136)<br>exon1:c.T1636C:p.S546P (591) |
| **Lipid metabolism** |  |  |  |
| *Aacs*, Chr 5, NM_030210 | exon8:c.770T>C:p.V257A (4799) | Longevity, health, limb clasping<br>***Atp8a1*↔*Aacs* Longevity, health** |  |
| *Apoa5*, Chr 9, NM_080434 | c.1061A>T:p.D354V (J_57L) | limb clasping, activity, muscle tone<br>***Col24a1*↔*Apoa5* weight**<br>***Hcn2*↔*Apoa5* muscle tone** |  |
| *Atp8a1*, Chr 5, NM_009727 | exon22:c.2037T>A:p.N679K (4799) | health, limb clasping, activity, muscle tone<br>***Atp8a1*↔*Aacs* Longevity, health** |  |
| *Cyp46a1*, Chr 12, NM_010010 |  |  | exon3:c.C263T:p.T88M (920_39R) |
| *Sqle*, Chr 15, NM_003129 | exon 6:c.1195C>T:p.R399X (895) | LOD 3.03 Longevity<br>***Rbbp8*↔*Sqle* Longevity** |  |
| *Tm7sf2*, Chr 19, NM_028454 | exon6:c.697T>C:p.Y233H (4654) | limb clasping, muscle tone, activity<br>***Dennd4a*↔*Tm7sf2* Longevity** |  |
| **Synapse function** |  |  |  |
| *Arhgef15*, Chr 11, NM_177566 | exon 2:c.275A>G:p.D92G (4751) | limb clasping, muscle tone, activity | exon2:c.A181T:p.I61F (A_230RR) |
| *Cd22*, Chr 7, NM_009845 | exon13:c.2414A>G:p.Q805R (520) | health, limb clasping<br>***Cd22*↔*Clint1* Longevity, muscle tone, activity** |  |
| *Celsr3*, Chr 9, NM_080437 | exon22:c.7415T>A:p.V2472E (918-15L) | Longevity, health, limb clasping | exon17:c.C6459A:p.C2153X (604)<br>exon4:c.T4604A:p.V1535E (A_347N) |
| *Dbnl*, Chr 11, NM_001146308 | exon8:c.709T>A:p.S237T (M199LL) | ns<br>***Birc6*↔*Dbnl* Longevity** |  |
| *Hcn2*, Chr 10, NM_008226 | exon 1:c.502C>T:p.R168C (J_57L) | Longevity, muscle tone, activity, weight<br>***Hcn2*↔*Apoa5* muscle tone** |  |
| *Kdm4a*, Chr 4, NM_001161823 | exon4:c.396T>A:p.Y132X (520) | Longevity, health, limb clasping |  |
| **Regulation of DNA activity** |  |  |  |
| *Gtf3c5*, Chr 2, NM_148928 | exon9:c.1169A>G:p.D390G (4654) | Longevity, muscle tone, activity |  |
| *Prdm15*, Chr 16 | splicing, 2099257392 A>G (352) | LOD 4.82 Longevity |  |
| *Zbtb41*, Chr 1, NM_172643 | exon3:c.1236C>A:p.N412K (M_333R) | Longevity | exon11:c.C2344A:p.Q782K (J_136N) |
| *Zzz3*, Chr 3 | chr3:152091574 C->T p.T to I (856) | LOD 3.61 Longevity |  |
| **Transcriptional repression** |  |  |  |
| *Ncor1*, Chr 11, NM_177229 |  |  | exon9:c.T866C:p.M289T (620) |
| *Ncor1*, Chr 11, NM_001252313 |  |  | exon17:c.A2043T:p.Q681H (S_165N) |
| *Rcor1*, Chr 12, NM_198023 |  |  | exon8:c.A650T:p.Q217L (4633) |
| *Sin3a*, Chr 9, NM_001110350 |  |  | exon13:c.C1907T:p.A636V (272) |
| *Tbl1xr1*, Chr 3, NM_030732 |  |  | exon12:c.G1081T:p.G361C (J_157N) |
| **Unknown** |  |  |  |
| *Clint1*, Chr11, NM_001045520 | exon12:c.1730T>C:p.M577T (520) | ns<br>***CD22*↔*Clint1* Longevity, muscle tone, activity** |  |
| *Col24a1*, Chr 3, NM_027770 | exon23:c.2542T>A:p.Y848N (J_57L) | ns<br>***Col24a1*↔*Apoa5* weight** |  |
| *Dennd4a*, Chr 9, NM_001162917 | exon15:c.1778C>T:p.A593V (4654) | ***Dennd4a*↔*Tm7sf2* Longevity** |  |

Animals were improved for longevity, additive subjective health score, limb clasping score, combined over three weeks, activity score, combined over three weeks, muscle tone score, combined over three weeks, or body weight at eight weeks. The tremor score did not change significantly for any lines, so is not included individually. The grooming score was an indicator of age and did not accurately reflect health, so is not included individually. ⇔ Indicates that pairwise interaction of candidate genes influenced the phenotype.

Four mutations (*Aacs, Apoa5, Atp8a1, Tm7sf2*) that were confirmed by mating and association results support previous publications that lipid metabolism is a primary pathway for pathogenesis (Buchovecky et al. 2013; Justice et al. 2013; Segatto et al. 2014; Kyle et al. 2016). The founding lesion in this pathway occurred in *Sqle*, a rate-limiting enzyme in cholesterol synthesis. MECP2 anchors a protein complex containing the nuclear receptor co-repressor 1 (NCOR1) to DNA, and mutations preventing MECP2’s interaction with the complex cause RTT (Ebert et al. 2013; Lyst et al. 2013). When MECP2 is absent, NCOR1 does not suppress its lipid synthesis targets, including *Sqle,* leading to lipid accumulation in the brain and liver of *Mecp2*-mutant males and females, resulting in metabolic syndrome (Buchovecky et al. 2013; Kyle et al. 2016). Acetoacetyl Co-A synthetase (*Aacs*) is another target of the NCOR1 co-repressor complex (Knutson et al. 2008), which allows ketone bodies to be used as an energy source (Hasegawa et al. 2012a; Hasegawa et al. 2012b).

Transmembrane 7 superfamily member 2 (*Tm7sf2*) functions just downstream of *Sqle* in the cholesterol biosynthesis pathway, and its function is linked to liver X receptor (LXR) and NFκβ signaling, suggesting roles in both lipid homeostasis and inflammation (Bellezza et al. 2013). Apolipoprotein A-V (*Apoa5*) is a key regulator of triglyceride metabolism (Gonzales et al. 2013). ATPase, aminophospholipid transporter (APLT), class I, type 8A, member (*Atp8a1*) is part of an ATPase-coupled membrane complex involved in vesicle trafficking and the transport of aminophospholipids (Paterson et al. 2006). Additional lesions were also observed that fall into the lipid metabolism pathway, including a mutation in line 920_39R that was not mated, which occurred in the neuron-specific cytochrome P450, family 46, subfamily a, polypeptide 1 (*Cyp46a1*). The product of CYP46A1 is 24*S*-OHC, a molecule essential for cholesterol turnover and a biomarker for abnormal cholesterol homeostasis in the *Mecp2* brain (Lund et al. 2003; Buchovecky et al. 2013).

A second group of mutations is predicted to affect synaptic function, as expected from previous studies of RTT patients and *Mecp2-*mutant mice (Chao et al. 2007; Shepherd and Katz 2011; Bellini et al. 2014). A mutation in cadherin, EGF LAG seven-pass G-type receptor 3 (*Celsr3*) was supported by mating and association analysis (*Celsr3^Sum15 m1Jus^*-p.V2472E), and two additional lines (604, A_347N) carried alleles. CELSR3 has a role in glutamatergic synapse formation and neuronal development (Feng et al. 2016; Thakar et al. 2017; Wang et al. 2017). Moreover, *Celsr3* is a target of the REST/Co-REST complex, which recruits other proteins including histone deacetylases (HDACs) and the transcriptional regulator SIN3a to mediate long-term gene silencing of target genes in neurons (Ballas et al. 2005b; Mandel et al. 2011; Monaghan et al. 2017; Hwang and Zukin 2018). Similarly, Drebrin-like (*Dbnl*) is a REST target involved in vesicular trafficking and synapse formation in neurons (Haeckel et al. 2008) whose candidacy is supported by mating and association analysis. Although we did not mate line 4633, it carries a missense mutation in co-REST, *Rcor1,* which binds and co-represses target genes with MECP2 (Ballas et al. 2005a). Other suppressor candidate genes that are predicted to function at the synapse are the hyperpolarization-activated, cyclic nucleotide-gated K+ 2 (*Hcn2*) and the Rho Guanine nucleotide exchange factor 15 (*Arhgef15*). Interestingly, *Hcn2* is a target of cAMP that regulates neuronal extension (Mobley et al. 2010) and mice that lack *Arhgef15* have increased excitatory synapse formation by allowing AMPA and NMDA receptors to be recruited to the synapse (Margolis et al. 2010). An additional allele of *Arhgef15* was observed in line A_230RR. CD22 antigen (*Cd22*) is a canonical B-cell receptor that clusters within the pathway of synaptic function because mutations in *Cd22* increase the ability of microglia to clear myelin debris and alpha-synuclein fibrils and remodel dendrites (Pluvinage et al. 2019).

The largest group of mutations affects DNA integrity or the DNA damage response (DDR). Three lines carried mutations in tet methylcytosine dioxygenase 1 (*Tet1*). TET1 modifies methylated CpG dinucleotides to 5-OH methylated bases in the first step of demethylation by base excision repair (BER) (Weber et al. 2016). *Tet1* plays a role in chromosomal stability and telomere length in embryonic stem cells through its methylation function (Yang et al. 2016). Although only one of the *Tet1* lesions (*Tet1^Sum18m1Jus^-*p.M81L) is supported by mating and association analysis, all three are missense mutations found in regions highly conserved throughout species, which lie outside the catalytic domain, but near the CpG binding domain (Supplemental Figure S1A). Mutations in the E2/E3 ubiquitin ligase baculoviral IAP repeat-containing 6 (*Birc6* also called BRUCE) were also found in three lines (Supplemental Figure S1B) and one was confirmed by mating and association analysis in line M_199L (*Birc6^Sum20 m1Jus^*-p.R3204Q). BIRC6 plays a role in stabilizing TRP53 in a first step towards inhibiting apoptosis (Ren et al. 2005). A non-inhibitor of apoptosis (IAP) function of BIRC6 is the regulation of the BRIT1-SWI-SNF double strand break (DSB)-response pathway (Ge et al. 2015). BIRC6-depleted cells display reduced homologous recombination repair, and *Birc6*-mutant mice exhibit repair defects and genomic instability (Lotz et al. 2004; Ren et al. 2005). However, BIRC6 has also recently been implicated in autophagy (Ebner et al. 2018). Spindlin 1 (*Spin1)* primarily plays a role in meiosis (Chew et al. 2013), while its induction in oocytes can trigger the DNA damage response; interestingly, the mutation lies within its third TUDOR-like domain (Supplemental Figure S2), which binds methylated histone H3, a mark associated with the DNA damage response (Choi et al. 2017; Shanle et al. 2017).

Additional mutations in this pathway are key to the repair of DSBs. A mutation in retinoblastoma binding protein 8 (*Rbbp8,* also called CtBP interacting protein *CtIP* or *Sae2* in yeast) was found in exon 15, a part of the critical C-terminal domain, which results in a premature stop codon that predicts a truncated protein (Figure 3A). RBBP8 is a determinant of DNA repair pathway choice (Escribano-Diaz et al. 2013). The presence of RBBP8 favors repair via the more error-free homologous recombination directed repair (HDR) over non-homologous end joining (NHEJ) during meiosis or in the S phase of dividing cells (Makharashvili et al. 2014). When RBBP8 is absent, NHEJ is the preferred choice for DDR repair. A survival curve shows that *Mecp2*/Y mice carrying the *Rbbp8* suppressor mutation demonstrate significantly improved longevity (Median survival of 112 days for *Mecp2*/Y*;Rbbp8*+/- versus 77 days for *Mecp2*/Y*;Rbbp8*+/+) (Figure 3B). Moreover, *Rbbp8* expression is increased in *Mecp2*/Y mouse brain at a symptomatic time point of 8 weeks (Figure 3C). Together, these data suggest that elevated *Rbbp8* in *Mecp2*-null mice may cause pathology that is ameliorated by reducing the amount of the protein.

**Figure 3:**
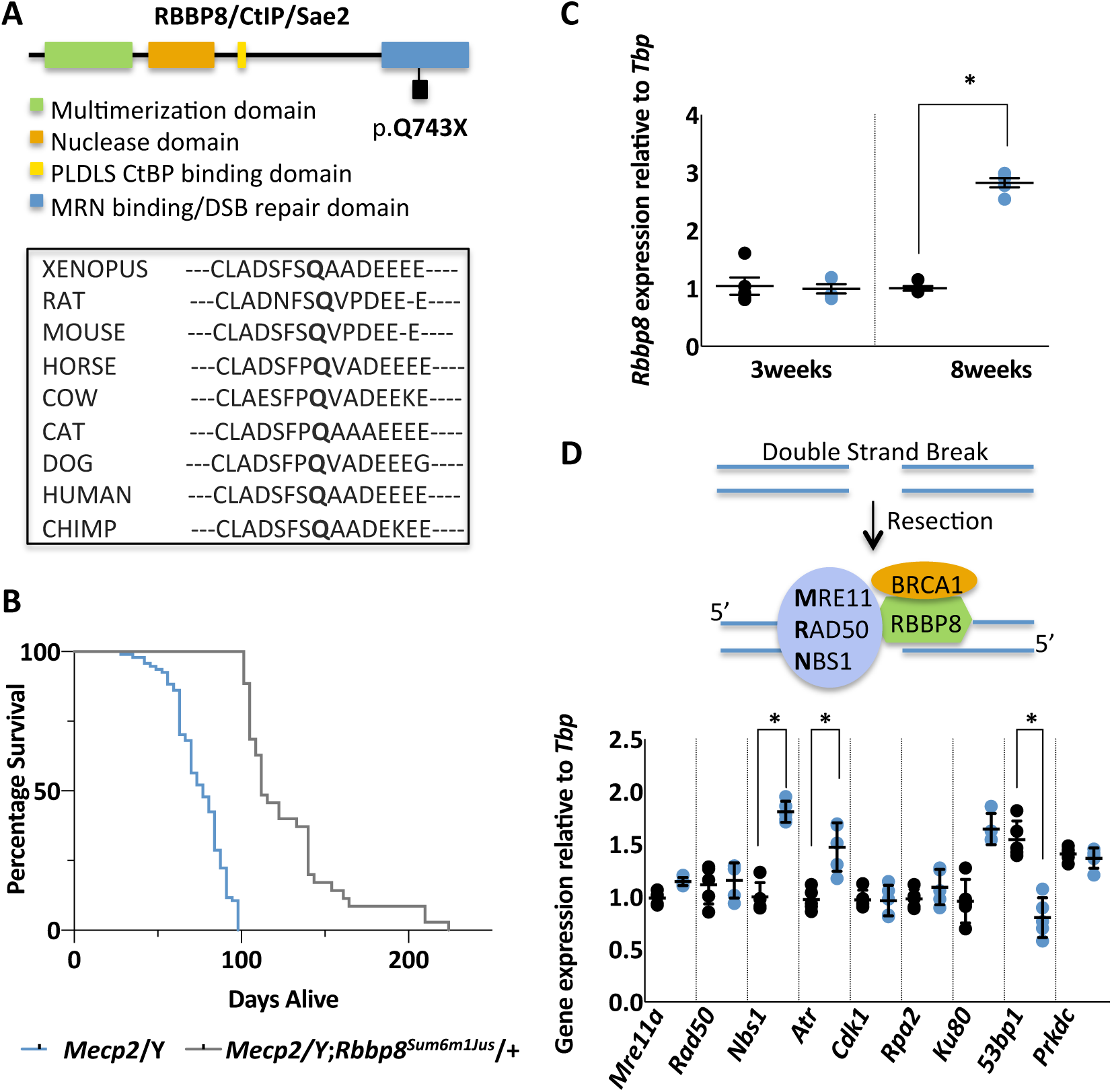
The DNA damage response (DDR) pathway is implicated in a *Mecp2/*Y mouse model of Rett syndrome. A. An ENU-induced nonsense mutation occurs at position p.Q743X within the critical C-terminal double strand break repair (DSB) domain (blue) of the mouse *Rbbp8* gene. Amino acid alignments generated using Clustal Omega show that this glutamine residue is well conserved among organisms from xenopus to human. B. *Mecp2/*Y*;Rbbp8^Sum6m1Jus^/*+ mice (gray) have increased longevity (n=35, Median survival 112 days) when compared to *Mecp2/*Y mice (blue) (n=94, Median survival 77 days) without secondary mutations (p<0.0001 by Mantel-Cox test). C. *Rbbp8* transcripts are elevated in symptomatic (8 week) *Mecp2/*Y (blue) brain compared to +/Y (black), but unchanged at 3 weeks. Results are representative of three independent experiments (p<0.01 by the two-sample Student’s t-test, +/Y: n=5, *Mecp2*/Y: n=5; error bars represent standard error of the mean (SEM). D. Double strand break repair by homologous recombination involves the recruitment of RBBP8 to the site of the break in an Mre11-Rad50-Nbs1 (MRN)/BRCA1-dependent manner. RBBP8 partners with BRCA1 to initiate DNA resection. Transcript levels of genes involved in the homologous recombination damage response (HDR) *Nbs1 and Atr* are elevated at 8 weeks in *Mecp2/*Y (blue) mice while *53bp1*, a gene involved in non-homologous recombination (NHEJ), is decreased compared to +/Y (black) (p<0.05 by the two sample Student’s t-test, +/Y: n=5, *Mecp2*/Y: n=5; error bars represent SEM).

Previous reports show that mesenchymal and neural stem cells derived from *Mecp2*/Y mice have an increased number of DSBs (Squillaro et al. 2010; Alessio et al. 2018). DSBs are initially detected by the Mre11-Rad50-Nbs1 (MRN) sensor complex, which activates the ATM or ATR kinases, leading to the phosphorylation of several downstream targets, including H2AX, which binds to the DSB (Falck et al. 2012). *Rbbp8* also plays a role in the removal of topoisomerase II (TOP2)-induced DNA DSBs, where it partners with BRCA1 to initiate DNA resection prior to repair (Aparicio et al. 2016) (Figure 3D). Many other factors are involved in the resolution of the DSB, including BRCA2 and FANCD2/FANCI-associated nuclease 1 (FAN1) (Orthwein et al. 2015). Interestingly, two mutations in *Brca1* and one in each of *Mre11a*, *Rad50*, *Fan1* and *Brca2* were also observed (Table 2, Supplemental Figure S2). To examine the extent of involvement of the DDR pathway, we examined the expression of several components of the HDR (*Mre11a, Rad50, Atr, Nbs1, Cdk1, Rpa2*), and NHEJ (*Ku80, 53bp1, Prkdc*) pathways in the brains of 8 week old *Mecp2*/Y and +/Y mice. Expression of Nijmegen breakage syndrome 1 (*Nbs1*), a member of the MRN complex, as well as the serine/threonine kinase ataxia telangiectasia and Rad3-related (*Atr)*, which is responsible for the transduction of signals from the MRN complex, were increased in the *Mecp2*/Y mouse brain. Conversely, the expression of p53 binding protein (*53bp1*) was decreased in *Mecp2*/Y brain (Figure 2D). 53BP1 functions in a manner antagonistic to RBBP8, blocking resection to promote NHEJ (Escribano-Diaz et al. 2013).

### Combinatorial mutations that improve health fall into different pathways

The investigation of pairwise gene-gene interactions in association analysis reveals six lines where the presence of two different mutations shows increased improvement in one or more traits (Figure 4A; Supplemental Table S2, S3). Five of the lines show improvement in longevity when two mutations are present: *Sqle* and *Rbbp8* in line 895, *Cd22* and *Clint1* in line 520, *Dennd4a* and *Tm7sf2* in line 4654, *Atp8a1* and *Aacs* in line 4799, and *Birc6* and *Dbnl* in line M_199LL. Two of the lines have interacting loci that do not show improved phenotypes alone, rather, they enhance the phenotype of another modifier locus: line 4654, (*Tm7sf2*óó *Dennd4a*; *Sum9i*), and line M_199LL, *Birc6*óó *Dbnl*; *Sum20i*). In line 520, the improvement of individual health traits segregated independently, suggesting two suppressors, so four N3 animals instead of two were sequenced (Supplemental Table S2). Subsequently, we found that mutations in both *Cd22* and *Kdm4a* independently improved traits, but did not further improve traits when inherited together. However, the combination of mutations in both *Cd22* and *Clint1* is also associated with improved activity and muscle tone, although *Clint1* did not improve traits when inherited alone. Similarly, a mutation in *Gtf3c5* in line 4654 improved longevity, activity and muscle tone independently of *Tm7sf2*. The mutation in *Tm7sf2* improved limb clasping, activity and muscle tone, yet longevity was not improved unless the animals also carried a mutation in *Dennda*. A further improvement in the health score is observed in line 4799 for animals that carried mutations in both *Atp8a1* and *Aacs*. In line J_57L, the combination of mutations in *Col24a1* and *Apoa5* is associated with lower body weight and in *Hcn2* and *Apoa5* with improved muscle tone. Notably, in most cases, the candidate genes belonged to two different pathways (Figure 4A). The exception is line 4799 where both *Aacs* and *Atp8a1* are predicted to function in lipid homeostasis, although in different aspects: utilization of energy and lipid trafficking, respectively. Thus, this combination has the potential to confer greater improvement together than either one alone.

**Figure 4:**
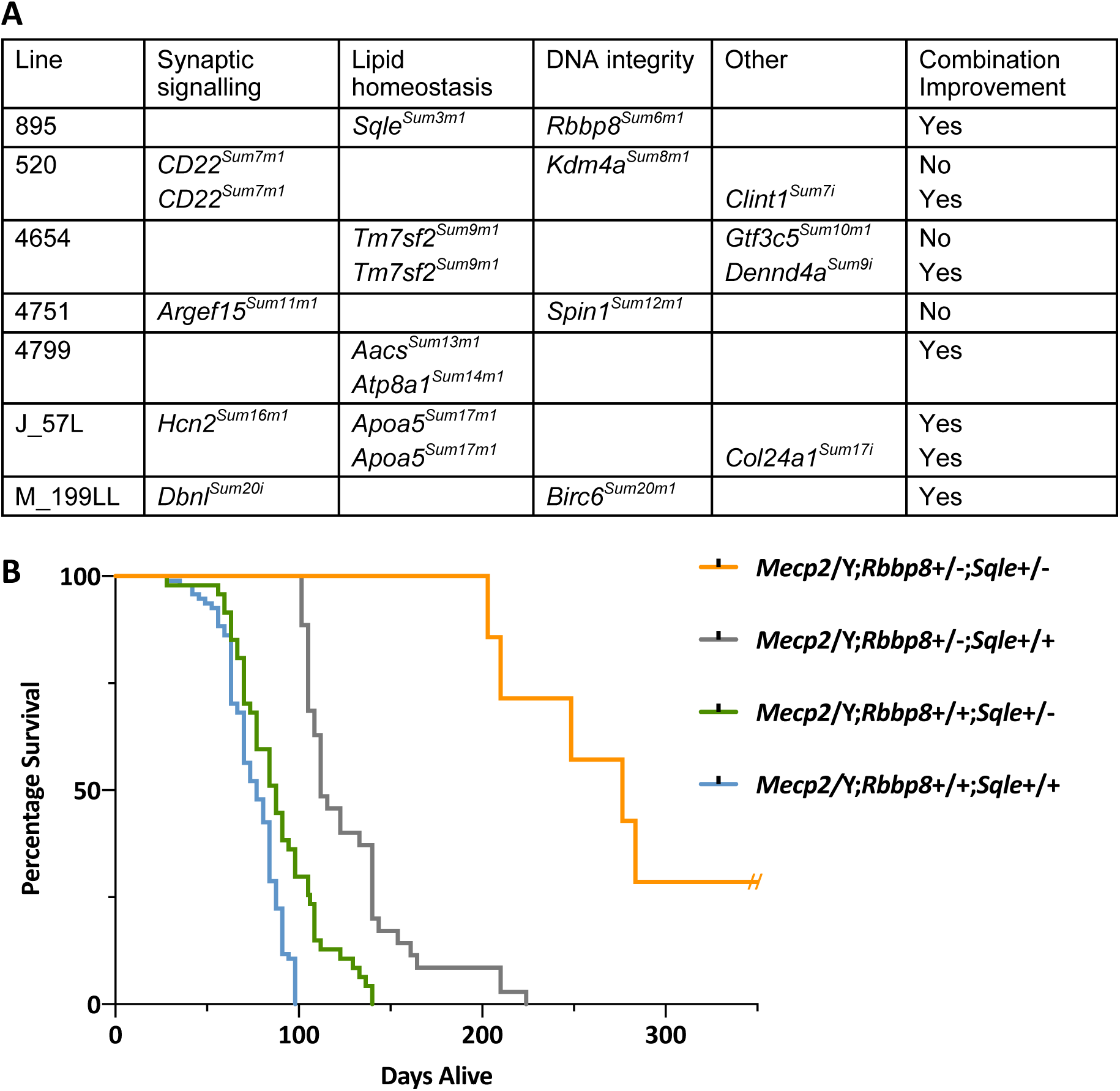
Combinatorial effects of multiple mutations on health and longevity A. Multiple lines (895, 520, 4654, 4751, 4799, J_57L and M_199LL) carry lesions in more than one gene that either independently or additively improve health. When two lesions are found in one line, the lesions generally affect two different pathways. With the exception of *Kdm4a* and *Cd22* in line 520, *Tm7sf2* and *Gtf3c5* in line 4654, and *Argef15* and *Spin1* in line 4751, the mutations all show positive combinatorial effects on *Mecp2/*Y health. B. *Mecp2/*Y mice from line 895, carrying mutations in *Sqle* and/or *Rbbp8,* show increased longevity when both mutations are present (*Mecp2/*Y;*Rbbp8+*/-;*Sqle+*/-) when compared to *Mecp2/*Y mice carrying either mutation alone (*Mecp2/*Y;*Rbbp8+*/-;*Sqle+*/+ or *Mecp2/*Y;*Rbbp8+*/+;*Sqle+*/-). *Mecp2/*Y;*Rbbp8+*/-;*Sqle+*/- n=7, Median survival 277 days; *Mecp2/*Y;*Rbbp8+*/-;*Sqle+*/+ n= 35, Median survival 112 days; *Mecp2/*Y;*Rbbp8+*/+;*Sqle+*/-, n= 47, Median survival 88 days; *Mecp2/*Y;*Rbbp8+*/+;*Sqle+*/+, n= 94, Median survival 77 days. (p<0.0001 by Mantel-Cox test, for each comparison of *Mecp2/*Y;*Rbbp8+*/+;*Sqle+*/+ to any of the other three groups). These data include the founder 895, N3 animals (genotyped for both *Rbbp8* and *Sqle*) and all of the N5 animals generated.

The founder of line 895 showed extreme longevity and was sacrificed at the age of 14 months. The pattern of inheritance of line 895 indicated that it carried two different suppressor mutations (Table 1; Supplemental Table S2). One, a nonsense mutation in *Sqle*, was identified by linkage (Chromosome 15, LOD 3.03), which was confirmed by the administration of statin drugs to mimic a downregulation of the cholesterol synthesis pathway (Buchovecky et al. 2013). N3 males from line 895 that had a longer life were bred to segregate and identify a second locus by backcrossing again to 129S6/SvEv animals to generate N4 and N5 generations (Supplemental Table S2). This analysis revealed that a nonsense mutation in *Rbbp8* (Chromosome 18) found in whole exome sequencing was consistent with trait improvement. Subsequent genotyping of animals from line 895 for *Sqle* and *Rbbp8* showed that although either mutation alone improved longevity (Median survival of 77 days for *Mecp2*/Y*;Rbbp8*+/+*;Sqle*+/+, 88 days for *Mecp2*/Y*;Rbbp8*+/+*;Sqle*+/-, 112 days for *Mecp2*/Y*;Rbbp8*+/-;*Sqle+/+*), the presence of both mutations conferred extreme longevity (Median survival of 277 days for *Mecp2*/Y*;Rbbp8*+/-*;Sqle*+/-), explaining a somewhat unusual survival curve in the initial report (Figure 4B (Buchovecky et al. 2013)). Seven animals from the N3 and N5 generations inherited both loci, and all lived longer than 29 weeks, with a range of 29 - 116 weeks (203 - 812 days). These data suggest that combining mutations from different pathways may further improve health in *Mecp2-null* mice.

## Discussion

Genetic modifier screens are mainly carried out in fruit flies, worms, yeast and bacteria to discover genes that are members of a developmental or biochemical pathway. The suppressor screen reported here represents the largest screen yet carried out in the mouse, and, by using massively parallel sequencing and association analysis, the screen identifies the largest number of candidate genes. Applications of modifier screens in the mouse are powerful (Carpinelli et al. 2004; Matera et al. 2008; Westrick et al. 2017). Even so, the mutations are identified based on their ability to suppress or enhance a mutant phenotype. Consequently, at least two mutations must be segregated to follow the phenotype, requiring an extensive amount of breeding. In the screen reported here, most lines showed evidence for complex trait inheritance, predicting that more than one modifier segregated, making mapping and candidate gene identification even more challenging. In a recent published modifier screen for thrombosis, very few candidate genes were identified, likely because of the lack of power in finding linkages using standard quantitative trait mapping strategies (Tomberg et al. 2018). Here, we employed statistical tests that assessed association instead of linkage to identify multiple interacting loci in six lines, four of which had an effect only in the presence of other modifiers (*Dbnl* with *Birc6*, *Dennd4a* with *Tm7sf2*, *Clint1* with *Cd22* and *Col24a1* with *Apoa5*).

ENU is expected to induce mutations randomly, but large genes are likely to have more mutations than small ones. *Ttn* is mutated in nearly every ENU screen because of its size (the cDNA is 81,843 base pairs, encompassing 192 exons), and twenty alleles were isolated here. Therefore, we cannot be certain that some genes had mutations because of their size, rather than their ability to modify *Mecp2*. *Tet1* had three alleles, and is a relatively small gene, which has 13 exons. *Birc6* also had three alleles, yet has 73 exons. Therefore, although alleles are called, some of the lesions may not be able to suppress phenotypes. This scenario demonstrates that mating all lines for inheritance and segregation is ideal. However, all of the founder lines reported here were not mated due to time, cost and animal space limitations. Our hope was that by identifying pathways, we could reduce the number of animals used in breeding. Candidate genes in any line can be confirmed after reanimation by IVF or by using CRISPR/Cas9 to engineer the mutation.

All of the potential modifying variants were not identified. As evidence, line 856 had a strong LOD score for a second modifying locus on Chromosome 6, yet WES did not identify a candidate in the region. Many factors can influence the observation that WES falls short of identifying all lesions. First, DNA quality can influence depth of coverage, making some heterozygous mutations difficult to call. Further, in previous ENU screens where sequence into introns was obtained, more lesions were found as potential splicing mutations (Justice et al. 1999; Boles et al. 2009), rather than the 3% reported here. Moreover, genome annotation in the mouse remains incomplete, with missing segments and incompletely annotated exons. For example, the suppressor in line 1395 mapped to a region of mouse Chromosome 7 that was not annotated, and therefore, was not present in the WES. Finally, loci present in the two inbred strains used here may also influence the penetrance of traits, yet modifying loci in the strain backgrounds were not a priority for identification in this study.

The inheritance of haplotypes can also make linked ENU-induced mutations difficult to sort as causative (Neuhaus and Beier 1998). In line A_333N, two mutations had very strong association scores as potential modifiers: a mutation in SH3 and multiple ankyrin repeat domains (*Shank1*) was a strong candidate gene based on its autism associations, while a mutation in *Fan1* would be consistent with the DDR pathway. However, these two genes lie only 20 Mb apart on Chromosome 7. Nine animals showed crossovers between the two loci, suggesting that the mutation in *Fan1* was the more strongly supported modifier, and there was no association with their inheritance together. In contrast, although *Aacs* and *Atp8a1* lie 60 Mb from each other on Chromosome 5, the mutations in each showed strong association with trait improvement, and their inheritance together further improved the traits. These data show that in these small family datasets without additional fine structure mapping crosses, candidate genes could be resolved by recombination and association analysis. However, the candidate genes are not confirmed by fine mapping, complementation or functional studies, so any should ideally be confirmed by additional experiments.

This unbiased modifier approach has identified key pathways in RTT pathogenesis, one of which supports the idea of improving synaptic signaling for treatment, as expected from multiple published evidences. Even so, our study suggests additional avenues for intervention. For example, CD22 responds to 2-hydroxypropyl-beta-cyclodextrin, which is in phase 2 and 3 clinical trials for Niemann-Pick disease, to reduce microglia-associated defects (Cougnoux et al. 2018), and antibodies directed against CD22 reduce microglial impairment in ageing brains (Pluvinage et al. 2019). In addition, two unknown pathways suggest that alternative pathways for intervention should be considered when studying RTT pathogenesis. The previously published pathway, lipid homeostasis, is supported by additional loci found here and by showing that lipid metabolism is directly regulated by an interaction between MECP2 and the NCOR1 co-repressor complex (Kyle et al. 2016). Mutations in *MECP2’s* NCOR interaction domain (NID) cause classical RTT in humans (Heckman et al. 2014) and RTT-like phenotypes in mice (Lyst et al. 2013), highlighting the importance of the NCOR1 interaction with MECP2 in RTT pathology. The NCOR complex is a master regulator of metabolism, playing roles in lipid biogenesis, glucose utilization and mitochondrial energy efficiency (Mottis et al. 2013). *Sqle* is a direct target of the MECP2-NCOR1-HDAC3 co-repressor function in the liver (Kyle et al. 2016). The data reported here show support for another target of this complex, *Aacs*, and another cholesterol synthesis enzyme *Tm7sf2*, as candidate suppressor genes.

This work suggests a role for DDR in RTT pathology for the first time. The genomic DNA of all cells must be protected from detrimental changes (Madabhushi et al. 2014) and MECP2’s absence is associated with the accumulation of genetic damage (Squillaro et al. 2010; Muotri et al. 2010). RBBP8 acts with BRCA1 to regulate the DSB repair choice between HDR or NHEJ (Huertas 2010), supporting the initial steps of DNA resection and recruiting other DNA repair proteins (Makharashvili et al. 2014). DSBs occur in all neural precursor cells and are important for normal neuronal development. However, the role of DNA integrity in mature neurons is less thoroughly understood, and is an active area of research. Recent studies suggest that DSBs arise in the brain as a by-product of neuronal activity (Suberbielle et al. 2013; Madabhushi et al. 2015). This in-turn impacts gene expression, especially of neuronal early response genes important for synapse development and maturation, neurite outgrowth, the balance between excitatory and inhibitory synapses and learning and memory. Robust mechanisms must be in place to rapidly and efficiently repair these DSBs; when these mechanisms fail, the accumulation of DSBs in the brain is a contributor to number of neurological diseases (Frappart and McKinnon 2008; Merlo et al. 2016). Normally, repair by HDR is more error free than NHEJ, and is thought to be more desirable. However, neurons may represent a unique case. DSBs are normally initiated by SPO11 in meiotic cells, which repair them by HDR with a homologous sister chromatid. In neurons that are transcriptionally active, topoisomerase II-mediated DSBs occur to relieve topological stress during transcription. Therefore, the elevated RBBP8 in *Mecp2-*null cells is most likely a result of increased DSBs; however, the repair of DSBs requires NHEJ in neurons, making HDR an undesirable event. Increased HDR and/or decreased NHEJ in *Mecp2-*null neurons may therefore be key to understanding RTT pathology. Modulating RBBP8 expression using the drug triapine has been an effective treatment to reduce HDR in cancer cells (Lin et al. 2014).

Altogether, the suppressor mutations implicated thus far paint a picture of altered metabolism and DNA damage that modulate synaptic function to cause pathology in Rett syndrome. Interestingly, many of the lines with the largest degree of symptom improvement carry at least two modifiers. Furthermore, combining modifiers from two different pathways improves symptoms, suggesting that combination therapies for RTT will be more effective than any single therapy. The results underscore the power of a genetic screen for understanding RTT biology as they demonstrate how a modifier screen in mammals is possible, especially for disease genes that are not present in more tractable genetic organisms. With new sequencing technologies and statistical approaches, such a screen should be amenable for nearly any gene for which phenotypes can be clearly and quantitatively assessed. Modifier screens in model organisms may thus help to identify the multitude of genetic variants that influence human disease presentation (Enikanolaiye and Justice 2019). Moreover, a genetic approach to disease prevention could revolutionize translational biology in neurological disorders.

## Methods

### Animals

All animal experiments were conducted under protocols approved by local Animal Care and Use Committees at Baylor College of Medicine (BCM) or at The Centre for Phenogenomics (TCP) accredited by the American Association for Laboratory Animal Care (AALAC) and Canadian Council on Animal Care (CCAC), respectively. Congenic 129.*Mecp2^tm1.1Bird^*/+ female mice were maintained by backcrossing females to males of the 129S6/SvEvTac strain. 129.*Mecp2^tm1.1Bird^*/Y (*Mecp2*/Y, which are *Mecp2*-null) and age matched wild type (+/Y) littermate controls were housed in plastic Techniplast cages with corncob bedding in rooms alternating 12-hr and 12-hr periods of light and dark, were provided acidified water and a Harlan Teklad 2920X diet *ad libitum* (19.1% protein, 6.5% fat; 0% cholesterol) (BCM) or Harlan Teklad 2919 (19% protein, 9% fat) (TCP). C57BL/6J males were obtained from The Jackson Laboratory (Bar Harbor, ME) at six weeks of age, and injected with three weekly doses of 100 mg/kg ENU at eight weeks as described (Kile et al. 2003). After recovery of fertility, ENU-treated males were mated to 129.*Mecp2^tm1.1Bird^/+* females, and their N1 male offspring were genotyped for the *Mecp2* mutation according to The Jackson Laboratory standard protocol using an Applied Biosystems thermocycler and resolution on a QiAxcel (Qiagen, Inc).

For thirty of the founder lines, N1 males showing signs of improvement were mated to 129S6/SvEvTac females or sperm was frozen and in vitro fertilization (IVF) was performed with oocytes from 129S6/SvEvTac females by the Cryopreservation and Recovery Core, The Centre for Phenogenomics. The N2 offspring from natural matings or from IVF were mated to 129.*Mecp2^tm1.1Bird^*/+ females or 129S6/SvEvTac males to produce N3 animals for inheritance testing and linkage or association analysis.

### Sequencing pipeline

DNA was extracted from mouse tails using standard methods. Briefly, tails were digested overnight at 55°C in tissue lysis buffer containing proteinase K, followed by purification with phenol chloroform, and precipitation with isopropanol. Resuspended DNA pellets were quantified using both a Nanodrop (Thermo Fisher) and Qubit (Thermo Fisher). For sequencing by The Centre for Applied Genomics, 500 ng of genomic DNA was fragmented to 200-bp on average using a Covaris LE220 instrument. Sheared DNA was end-repaired and the 3’ ends adenylated prior to ligation of adapters with overhang-T. Genomic libraries were amplified by PCR using 10 cycles and 750 ng of the PCR-enriched library was hybridized with biotinylated probes that target exonic regions; the captured exome libraries were amplified by an additional 10 cycles of PCR. Exomic libraries were validated on a Bioanalyzer 2100 DNA High Sensitivity chip (Agilent Technologies) for size and by qPCR using the Kapa Library Quantification Illumina/ABI Prism Kit protocol (KAPA Biosystems) for quantities. Exome libraries were pooled and sequenced with the TruSeq SBS sequencing chemistry using a V4 high throughput flowcell on a HiSeq 2500 platform following Illumina’s recommended protocol. Approximately 6-8 gigabases of raw paired end data of 126-bases were generated per exome library.

Sequence reads were mapped to the mm10 reference genome with *bwa* (Li and Durbin 2010), and variants were called with *samtools* (Li et al. 2009) requiring a minimum SNP quality of 20 and a read depth ≥ 5. Putative Insertion-Deletion (InDel) calls were realigned and SNP quality was recalibrated with GATK. Custom scripts were used to annotate variants with respect to their predicted impact on protein sequence. To remove inbred strain polymorphisms as well as systematic sequencing artifacts, variants were removed from consideration if identified in the parental strains, dbSNP, or other founder males sequenced in this study.

Candidate lesions were confirmed by Sanger sequencing prior to genotyping N3 animals. Primer sequences are available upon request.

### Quantitative Reverse Transcription Polymerase Chain Reaction (qRT-PCR)

Whole brain tissue was homogenized using a 5mm stainless bead and Qiagen Tissue Lyser II in QIAzol Reagent according to manufacturer’s instructions. Total RNA was then isolated using the RNeasy Lipid Tissue Mini Kit (Qiagen). cDNA was prepared by reverse transcription using the Superscript VILO cDNA synthesis kit (Invitrogen). Quantitative reverse transcription PCR (qRT-PCR) was performed in triplicate for each sample on a Viia7 instrument (ABI) using Power SYBR Green PCR Master Mix (Invitrogen). The PCR conditions were as follows: 95°C for 10 min, 40 cycles of 95°C for 15 s and 60°C for 60 s. Gene expression was normalized to TATA-binding protein (*Tbp*) as an internal loading control and results analyzed using the 2^−(ΔΔCT)^ method. Gene expression was displayed as the expression in *Mecp2/*Y mice relative to that of age matched +/Y littermate controls. Primer sequences are available upon request.

### Statistical Methods

Complex versus Mendelian inheritance was assessed from the number of animals that inherited trait improvement or longevity in each N2 family (Supplemental Table S2). Column 4 of Table 1 reports the p-value of the Fisher’s exact test for testing the Mendelian hypothesis calculated based on 10^6^ Monte-Carlo simulations. For each line, the observed decomposition of the N3 improved cases across N2 families is tested for the Mendelian hypothesis by generating data conditional on whether or not N2 animals inherited the trait given the presence (with probability 0.50) or absence (with probability 0.25) of an N3 offspring with improvement in the family. The p-value is then calculated considering the number of as-or-more extreme cases under the same family configuration than the one observed.

In screen 2, a standard quantitative trait loci (QTL) analysis was not powerful enough to identify modifier loci because genome-wide SNP analysis for linkage was not carried out, and each N2 family had a relatively low number of animals. To investigate associations of genetic loci with outcomes based on subjective health parameters, we used the cumulative link models (CLM) suitable for ordinal data (Agresti 2011; Christensen 2015). Specifically, we considered the Health Score (h) as well as the limb clasping (lc), tone (mt), activity (a) and tremor (t) scores, and fitted separate models to each. The genetic associations with the body weight (nominal value at the time of sacrifice) were inferred using linear models. In both CLM and linear models, random effects were included to account for potential clustering of outcomes within families since mice from the same family are expected to show similar characteristics. However, in almost all cases, random effects had a negligible variance suggesting that there was no significant clustering of the outcomes, allowing for the fitted models to include only fixed effects. For each line, we investigated marginal effects of each gene, as well as pairwise gene-gene interactions on these traits. The associations of genetic loci with the lifetime (time to sacrifice) of mice were assessed using parametric survival models, where the final model for each line was selected using the Akaike Information Criterion (AIC) (Akaike 1974). Similarly, we assessed the marginal effect of each gene and pairwise gene-gene interactions in these survival models. All association analyses were performed using the R statistical programing software (R Core Team 2013), along with the packages ordinal (Akaike 1974), lme4 (Bates et al. 2015), and flexsurv (Jackson 2016). Kaplan-Meier survival curves in Figures 2B and 4B were generated in GraphPad Prism 8 followed by the Mantel-cox test (log-rank comparison). Comparisons between two groups, as in qPCR experiments were carried out in GraphPad Prism 8 using the two-sample Student’s t-test.

## Acknowledgements

The Centre for Applied Genomics (TCAG) generated sequence for founder animals from screen 2. The Canadian Mutant Mouse Repository Cryopreservation and Recovery Core at the Centre for Phenogenomics (TCP) carried out the sperm freezing and *in vitro* fertilization. We thank Dr. Dan Durocher for advice in examining the DDR pathway, Ms. Ashlee Dargie, Mr. Travis Brooke-Bisschop and Mr. Stephen McDonald for technical assistance, and Dr. Rebekah Tillotson and Ms. Neeti Vashi for critical review. This work was supported by grants from the Rett Syndrome Research Trust and the Canadian Institute for Health Research Foundation Scheme File# 375397 to MJJ.

## Author contributions and disclosure declarations

MJJ conceived of the work, obtained funding and wrote the manuscript. AE generated data for DSB pathway, performed functional network analysis and wrote the manuscript. JR oversaw the screen, organized data, and wrote the manuscript. CMB, CT, and MS generated data. SP, RZ and JS generated and analyzed sequencing data. EA performed statistical analysis and wrote the manuscript.

